# Colistin resistance mutations in *phoQ* sensitize *Klebsiella pneumoniae* to IgM-mediated complement killing

**DOI:** 10.1101/2022.09.15.508115

**Authors:** Sjors P.A. van der Lans, Manon Janet-Maitre, Frerich M. Masson, Kimberly A. Walker, Dennis J. Doorduijn, Axel B. Janssen, Willem van Schaik, Ina Attrée, Suzan H. M. Rooijakkers, Bart W. Bardoel

**Affiliations:** Department of Medical Microbiology, University Medical Centre Utrecht, The Netherlands; Bacterial Pathogenesis and Cellular Responses group, UMR5075, Institute of Structural Biology, University Grenoble Alpes, UMR5075 France; Department of Microbiology and Immunology, University of North Carolina, USA; Department of Fundamental Microbiology, University of Lausanne, Switzerland; Institute of Microbiology and Infection, College of Medical and Dental Sciences, University of Birmingham, United Kingdom

## Abstract

The Gram-negative bacterium *Klebsiella pneumoniae* is notorious for a strong increase of infections with antibiotic resistant strains. To treat infections with antibiotic resistant *K. pneumoniae,* clinicians increasingly need to use the last resort antibiotic colistin. *K. pneumoniae* can develop colistin resistance by modifying its membranes. During infection the membranes of Gram-negative bacteria are also targeted by the human immune system via the complement system. Gram-negative bacteria have an outer and inner membrane separated by a thin cell wall. Activation of the complement system leads to the formation of the membrane attack complex (MAC), a pore that inserts into the outer membrane, and ultimately leads to lysis of the bacterium. As both colistin and the MAC interact with the outer membrane of Gram-negative bacteria, we wondered if developing colistin resistance influences MAC-mediated killing of *K. pneumoniae*.

Using clinical isolates that developed colistin resistance, we found that the strain Kp209_CSTR became more sensitive to MAC-mediated killing compared to the wild-type strain. MAC-mediated membrane permeabilization of Kp209_CSTR required antibody dependent activation of the classical complement pathway. Strikingly, Kp209_CSTR was bound by IgM in human serum that did not recognise the wild-type strain. Depletion of Kp209_CSTR-specific antibodies from serum prevented MAC-mediated membrane permeabilization, which was restored by adding back IgM. Genomic sequence comparison revealed that Kp209_CSTR has a deletion in the *phoQ* gene. RNAseq analysis suggested that this mutation locks PhoQ in a constitutively active state. These results indicate that PhoQ activation in Kp209_CSTR leads to both colistin resistance and sensitivity to MAC-mediated killing. Together, our results show that developing colistin resistance can sensitize *K. pneumoniae* to killing by the immune system.

## Introduction

Infections with antibiotic-resistant bacteria form a serious threat for public health. One of the most dramatic rises in multi drug-resistance has been seen for infections with the Gram-negative bacterium *Klebsiella pneumoniae* (Bush et al., 2011; Laxminarayan et al., 2013). Infections with *K. pneumoniae* caused nearly 1 million antibiotic resistance associated deaths in 2019 alone (Murray et al., 2022). To treat infections with antibiotic resistant bacteria, clinicians increasingly need to use last resort antibiotics such as colistin. Colistin, a membrane-targeting antibiotic, is attracted to the Gram-negative cell envelope via electrostatic interactions (Andrade et al., 2020). The Gram-negative cell envelope consists of a thin peptidoglycan cell wall in between an outer and an inner membrane, with negatively charged lipopolysaccharides (LPS) being present in the outer leaflet of the outer membrane. The positively charged colistin is attracted to these negative charges in the cell envelope, causing colistin to insert into the membranes. This will subsequently destabilise and disrupt both the outer and inner membrane, ultimately leading to cell death (Sabnis et al., 2021). The use of colistin in the treatment of *K. pneumoniae* infections can lead to the development of colistin resistance in *K. pneumoniae* in clinical settings (Azam et al., 2021). Colistin resistance is mostly due to altered decoration of the lipid A part of LPS that reduces the negative electrostatic charge, such as addition of amino-4-deoxy-L-arabinose (L-Ara4N) and/or phosphoethanolamine (PEtn) (Sabnis et al., 2021). These LPS modifications can be caused by mutations in different genes, including those encoding the PmrAB and PhoPQ two-component systems, and the PhoPQ regulator MgrB (Bardet et al., 2017; Bray et al., 2022; Cannatelli et al., 2014; Gogry et al., 2021).

Development of colistin resistance in Gram-negative bacteria has been shown to influence both fitness and virulence (Choi et al., 2020; Janssen et al., 2021; López-Rojas et al., 2011). In addition, colistin resistance has been shown to induce resistance against cell envelope targeting compounds of the human immune system. For instance, *K. pneumoniae* that developed colistin resistance *in vitro* also developed resistance against the anti-microbial peptide LL-3 (Janssen et al., 2021). For *K. pneumoniae* colistin resistance has also been shown to lead to increased survival in human serum (Choi & Ko, 2015). But as serum contains many different antimicrobial compounds, it is not yet clear to which compounds the colistin resistant strains became less sensitive.

One of the main antimicrobial effectors in serum is the complement system, a protein network in serum and body fluids that can directly kill Gram-negative bacteria via the formation of the membrane attack complex (MAC). MAC formation is initiated after bacteria are recognized by the complement system, which leads to sequential deposition of complement proteins on the bacterial surface. This ultimately results in the formation of the MAC, a multi-protein complex that forms a pore in the outer membrane, leading to inner membrane destabilization and bacterial killing (Heesterbeek et al., 2019). *K. pneumoniae* can be resistant to killing by the MAC, for instance through the production of protective extracellular polysaccharides such as the capsule and O-antigen component of LPS (Jensen et al., 2020; Pennini et al., 2017). Furthermore, modifications of LPS and the O-antigen are known to affect MAC formation on *K. pneumoniae* (Doorduijn et al., 2021; Pennini et al., 2017). As LPS modifications affect both MAC formation and colistin resistance, it has been suggested that the LPS modifications mediating colistin resistance might affect MAC-mediated killing of *K. pneumoniae.*

In this study we investigated if colistin resistance affects MAC-mediated membrane permeabilization of *K. pneumoniae*, using clinical isolates that developed colistin resistance after *in vitro* evolution (Janssen et al., 2021). We found that one of the colistin resistant strains, Kp209_CSTR, became more sensitive to MAC-dependent membrane permeabilization via the classical complement pathway compared to the parental strain. Kp209_CSTR was bound by IgM present in human serum that did not recognise its colistin sensitive counterpart. We show that the Kp209_CSTR-specific triggers IgM MAC-mediated membrane permeabilization. Further analysis revealed that a mutation in *phoQ* of Kp209_CSTR locked PhoQ in a constitutively active state, suggesting that PhoQ activation results in both colistin resistance and increased MAC-mediated killing. Together, our results show that PhoQ activity has a opposite effect on colistin resistance and MAC-mediated membrane permeabilization in *K. pneumoniae*.

## Results

### Inner membrane damage correlates with MAC-mediated killing of *K. pneumoniae*

We previously showed that inner membrane damage correlates with bacterial killing of Gram-negative bacteria using a fluorescent DNA dye (SYTOX) that can only stain Gram-negative bacteria when both the outer and inner membrane are damaged (Heesterbeek et al., 2019). To determine if MAC-induced membrane permeabilization also correlates with killing of *K. pneumoniae* clinical isolates, we analysed the MAC-mediated membrane permeabilization and killing of ten clinical *K. pneumoniae* strains. Bacteria were incubated with 10% normal human serum (NHS) containing both complement and naturally occurring antibodies. Inner membrane permeabilization was monitored by measuring fluorescence over time and compared to bacterial survival that was assessed by counting colony forming unites (CFU) on plate (fig. 1ab).

**Figure 1.**
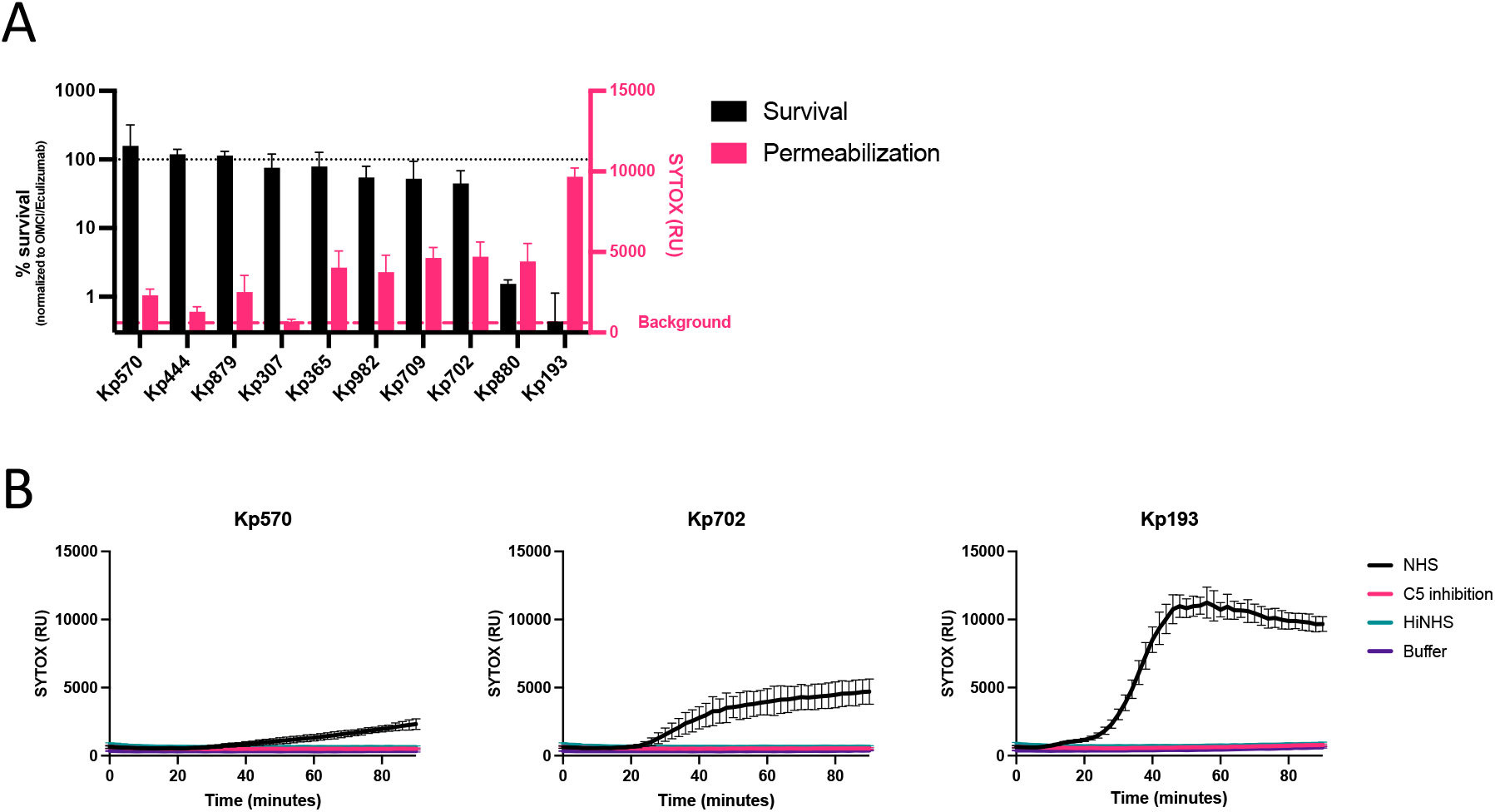
MAC-dependent inner membrane permeabilization correlates with reduced viability of *K. pneumoniae*. (A) Survival on plate and inner membrane permeabilization of clinical *K. pneumoniae* isolates after 90-minute incubation in 10% normal human serum (NHS) at 37°C in the presence of 1 μM SYTOX green nucleic acid stain. Survival data was normalized to CFU counts in conditions where C5 conversion was inhibited by addition of 20 μg/ml OMCI and 20 μg/ml Eculizumab. Inner membrane permeabilization (SYTOX fluorescence intensity) was determined in a microplate reader. Red dotted line indicates background (OMCI+ Eculizumab). (B) Inner membrane permeabilization of Kp570, Kp702 and Kp193 in the presence of 10% NHS, 10% NHS in which C5 conversion was inhibited by addition of 20 μg/ml OMCI and 20 μg/ml Eculizumab (C5 inhibition), or 10% heat inactivated NHS (HiNHS). Bacteria were incubated at 37°C in the presence of 1 μM SYTOX green nucleic acid stain, and inner membrane permeabilization (SYTOX fluorescence intensity) was detected every 2 minutes for 90 minutes in a microplate reader. (A-B) Data represent mean ± standard deviation of three independent experiments.

First, we observed that three out of ten tested isolates were resistant to MAC-mediated killing (Kp570, Kp444 and Kp879), as these strains showed no decrease in survival (fig. 1a). We observed no or a minimal increase in inner membrane damage for these three MAC resistant isolates (fig. 1ab, S1). Furthermore, the strain that was most sensitive to MAC-mediated killing (Kp193) showed the strongest membrane permeabilization (fig. 1a). Inner membrane damage started after 20 minutes in Kp193, after which the signal quickly increased and reached a plateau after 40 minutes (fig. 1b). Five strains showed a slow increase in membrane permeabilization (Kp365, Kp982, Kp709, Kp702 and Kp880) (fig. 1ab, S1). Most of these strains showed only a minor reduction in survival, except for Kp880. When complement activation was abrogated by heat-inactivation of NHS, or when MAC formation was blocked by inhibition of C5 conversion, no inner membrane permeabilization was observed (fig. 1b, S1). Taken together, these data indicate that MAC-dependent inner membrane damage correlates with killing of clinical *K. pneumoniae* strains.

### Developing colistin resistance sensitizes *K. pneumoniae* Kp209 to MAC-mediated killing

To study the influence of developing colistin resistance on MAC-mediated killing of *K. pneumoniae*, we used isogenic *K. pneumoniae* strain pairs that were previously selected during an *in vitro* evolution experiment (Janssen et al., 2021). Herein, three colistin sensitive *K. pneumoniae* strains (Kp209, Kp257 and Kp040) were exposed to increasing concentrations of colistin to evolve colistin resistance. All colistin-resistant (CSTR) strains (Kp209_CSTR, Kp257_CSTR and Kp040_CSTR) had altered Lipid A that reduced its negative charge (Janssen et al., 2021).

The three isogenic strain pairs were incubated with NHS and MAC-mediated membrane damage was determined. While acquisition of colistin resistance did not affect the MAC-dependent permeabilization of Kp257 and Kp040, we observed that Kp209_CSTR was more sensitive to MAC-mediated membrane permeabilization compared to its parental strain (fig. 2a). Membrane damage of the colistin-resistant strain was caused by MAC, as this effect was blocked by heat-inactivation of NHS or the inhibition of C5 conversion. In concordance, only 1% survival was observed for Kp209_CSTR in 10% NHS (fig. 2b). In contrast, the wild-type Kp209 showed not decrease in survival (fig. 2b). Together, these data suggest that developing colistin resistance sensitised Kp209 to MAC-mediated killing in NHS.

**Figure 2.**
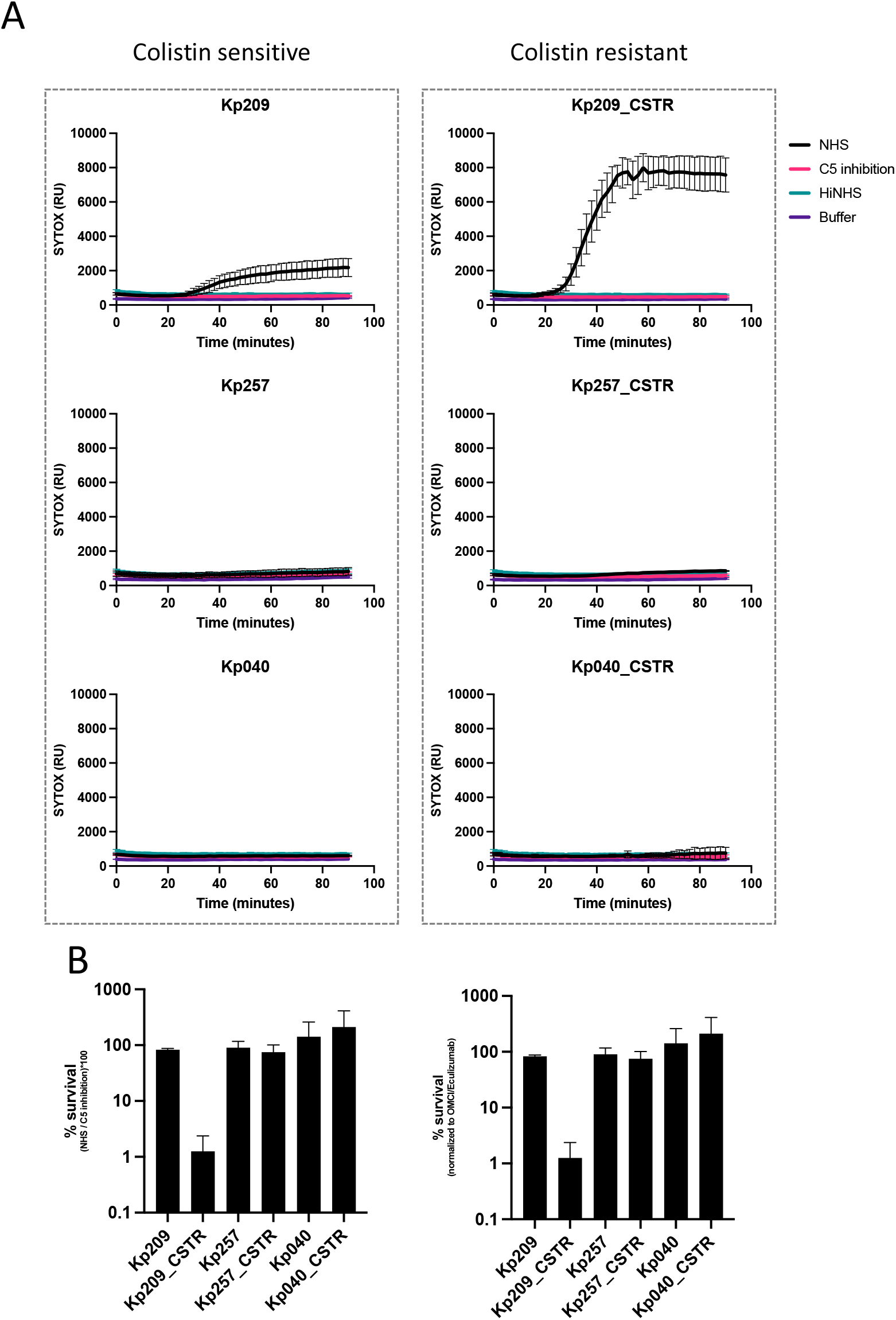
Colistin resistant Kp209_CSTR has been sensitized to MAC-mediated killing. (A) Inner membrane permeabilization of Kp209, Kp257 and Kp040, and their colistin resistant derived strains Kp209_CSTR, Kp257_CSTR and Kp040_CSTR in the presences of 10% NHS, 10% NHS in which C5 conversion was inhibited by addition of 20 μg/ml OMCI and 20 μg/ml Eculizumab (C5 inhibition), or 10% heat inactivated NHS (HiNHS). Bacteria were incubated at 37°C in the presence of 1 μM SYTOX green nucleic acid stain, and inner membrane permeabilization (SYTOX fluorescence intensity) was detected every 2 minutes for 90 minutes in a microplate reader. (B) Survival on plate of Kp209, Kp257 and Kp040, and their colistin resistant derived strains Kp209_CSTR, Kp257_CSTR, and Kp040_CSTR after 90 minutes incubation in 10% NHS at 37°C. Survival data is normalized to CFU counts in conditions where the terminal complement pathway was blocked by inhibiting C5 conversion (20 μg/ml OMCI and 20 μg/ml Eculizumab). (A-B) Data represent mean ± standard deviation of three independent experiments.

### Classical pathway activation is crucial for killing of Kp209_CSTR

To study the mechanism that caused increased MAC-mediated killing of Kp209_CSTR, we first analysed which complement pathway was responsible for MAC-dependent permeabilization. The complement system can be activated via three distinct pathways (classical, lectin and alternative pathway) which are activated via different mechanisms, but all lead to the formation of MAC on Gram-negative bacteria. Both the classical and the lectin pathway depend on complement protein C2, therefore we used C2 depleted serum to determine the involvement of these pathways. We observed that depletion of C2 resulted in the complete loss of membrane permeabilization of Kp209_CSTR (fig. 3a). This effect was restored by adding back C2 to physiological levels, which indicated that C2 was crucial for complement-mediated membrane damage of Kp209_CSTR. Similarly, factor B (fB) depleted serum was used to determine the role of the alternative pathway. Depletion of fB did not influence inner membrane permeabilization of Kp209_CSTR (fig. 3b). These data suggest that killing of Kp209_CSTR is primarily driven by the classical and/or lectin pathway. To further distinguish between the classical and lectin pathway, we used inhibitors to specifically block the first steps of classical pathway activation. The initiation of the classical pathway starts when IgG or IgM antibodies bind the bacterial surface. Upon binding, these antibodies can recruit the large C1 complex, which is specific to the classical pathway. C1 consists of the recognition protein C1q and associated proteases C1r and C1s. To block C1 dependent complement activation, we simultaneously added both a monoclonal antibody that directly prevents C1q association to surface-bound antibodies, and the C1r inhibitor BBK32 (Garcia et al., 2016; McGonigal et al., 2016; Zwarthoff et al., 2021). Addition of these inhibitors to NHS prevented inner membrane damage of Kp209_CSTR (fig. 3c). In summary, these findings indicate that MAC-mediated membrane permeabilization of Kp209_CSTR in these conditions primarily depends on the classical pathway.

**Figure 3.**
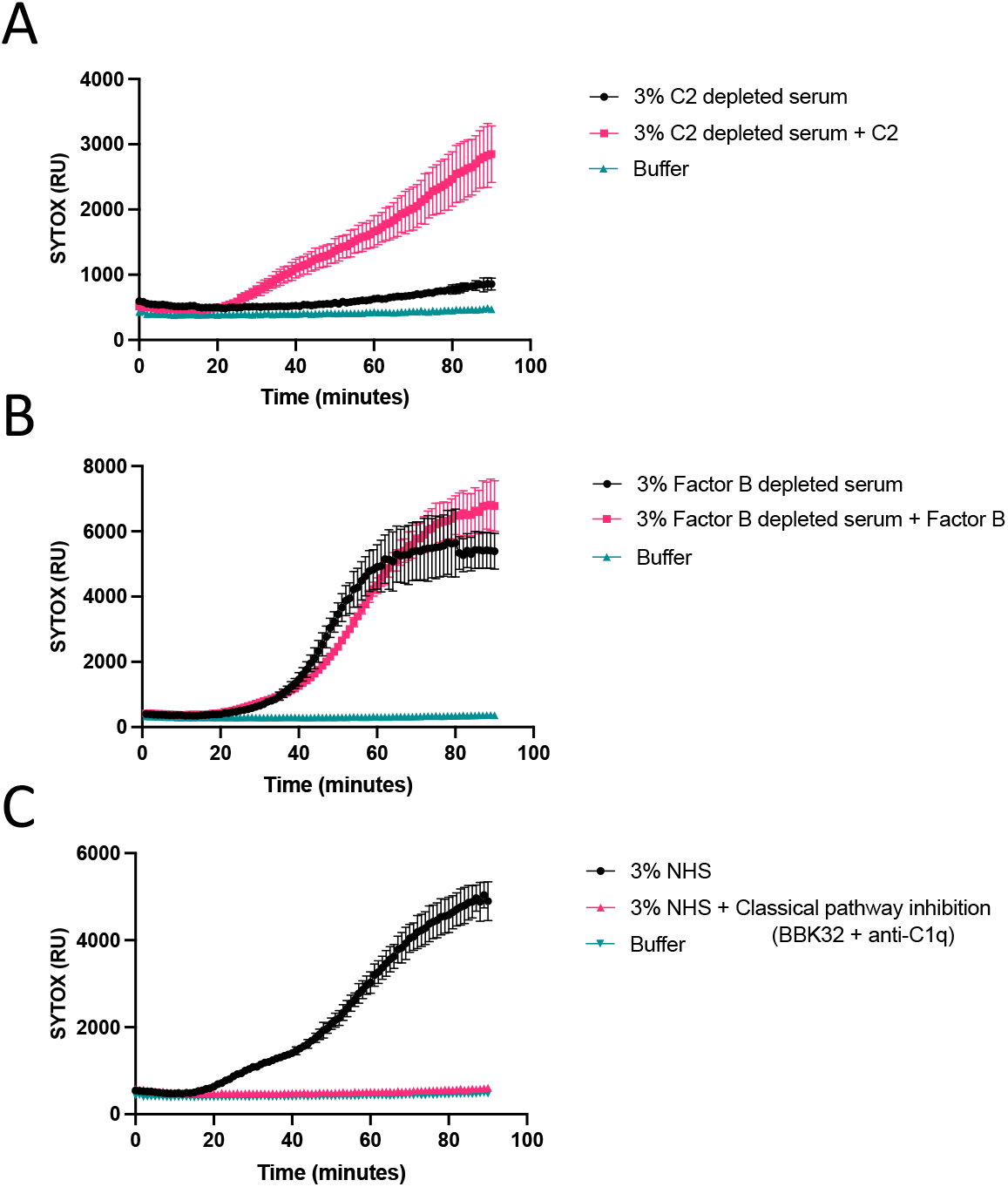
MAC-dependent inner membrane permeabilization of Kp209_CSTR is dependent on the classical pathway. (A) Inner membrane permeabilization of Kp209_CSTR in the presence of 3% C2 depleted serum or C2 depleted serum reconstituted with C2 to physiological concentrations (0.6 μg/ml in 3% serum). (B) Inner membrane permeabilization of Kp209_CSTR in the presence of 3% fB depleted serum or fB depleted serum reconstituted with fB to the physiological concentration (6 μg/ml in 3%). (C) Inner membrane permeabilization of Kp209_CSTR in the absence or presence of the classical pathway inhibitors anti-hu-C1q and BBK32 (both 10 μg/ml) in 3% normal human serum (NHS). (A-C) Bacteria were incubated at 37°C in the presence of 1 μM SYTOX green nucleic acid stain, and inner membrane permeabilization (SYTOX fluorescence intensity) was detected every two minutes for 90 minutes in a microplate reader. Data represent mean ± standard deviation of three independent experiments.

### IgM specific for Kp209_CSTR is responsible for MAC-mediated membrane permeabilization of Kp209_CSTR

The sensitivity of Kp209_CSTR to the classical complement pathway suggests a role for IgG or IgM in the increased MAC-dependent membrane permeabilization in NHS. We hypothesized that there might be increased binding of IgG or IgM to Kp209_CSTR in NHS. Incubation of Kp209 and Kp209_CSTR with NHS revealed that there was no major difference in total IgG (fig. S2a) or IgM (fig. S2b) binding to these strains. As IgG and IgM in NHS are polyclonal, we analysed if NHS contains antibodies that specifically recognize Kp209_CSTR, but not Kp209. A relative low concentration of Kp209_CSTR-specific antibodies may explain why there is no difference observed in the total antibody binding between Kp209 and Kp209_CSTR. To investigate this hypothesis, we performed antibody depletion using whole bacteria to test whether NHS contained Kp209_CSTR-specific antibodies (fig. 4a). NHS was incubated with Kp209, Kp209_CSTR, or *E. coli* used as a control, at 4°C to allow antibody binding without activating complement, followed by the removal of bacteria to collect NHS deprived of bacterium-specific antibodies. Three rounds of depletion where sufficient to remove most strain-specific antibodies (fig. 4b & fig. S2c). Depleting NHS with Kp209 removed all IgG binding to Kp209_CSTR (fig. 4b), and *vice versa* (fig. S2c), indicating that both strains were bound by the same IgG’s in NHS. Depletion with Kp209_CSTR abolished IgM binding to Kp209 (fig. S2c), showing that all the IgM that bound Kp209 also recognized Kp209_CSTR. However, although depletion with Kp209 strongly reduced IgM binding to Kp209_CSTR, considerable binding was still detectable, implying that NHS contains Kp209_CSTR-specific IgM (fig. 4b & fig. S2e). We validated that antibodies were specifically removed from NHS by *K. pneumoniae* depletion, as they did not affect antibody binding to *E. coli* (fig. S2d).

**Figure 4.**
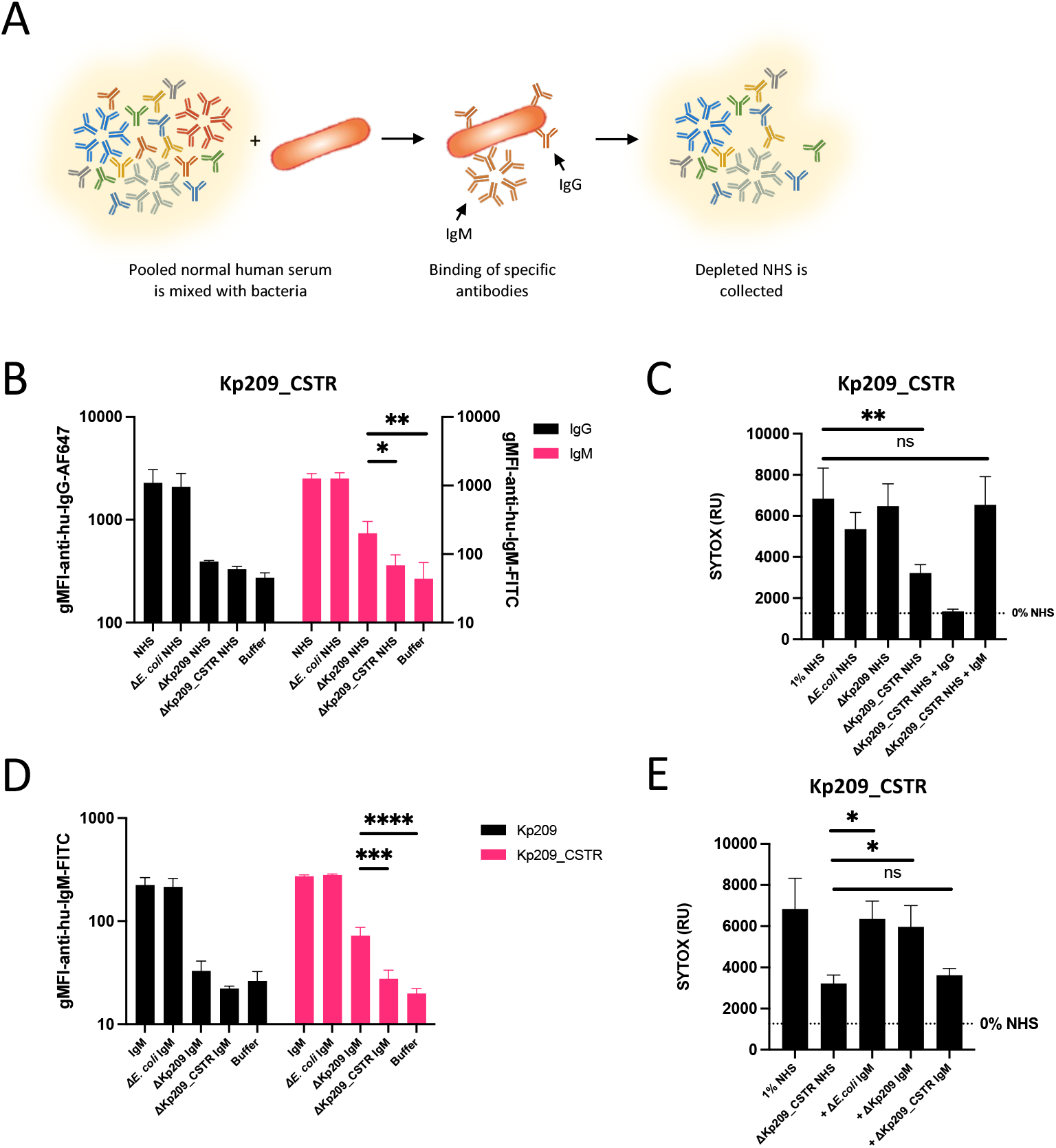
NHS contains Kp209_CSTR specific IgM that is vital for MAC-dependent inner membrane permeabilization. (A) Schematic representation of method to deplete bacterium specific antibodies from normal human serum (NHS). NHS was mixed with bacteria and incubated for 10 minutes at 4°C to allow specific antibodies to bind. Bacteria were pelleted and discarded, and the depleted NHS was collected. Three rounds of depletion were performed. (B) Kp209_CSTR was incubated with NHS, NHS depleted with *E. coli* MG1655, Kp209 or Kp209_CSTR (Δ*E. coli* NHS, ΔKp209 NHS, and ΔKp209_CSTR NHS, respectively) for 30 minutes at 4°C. Binding of IgG was measured in 0.3% NHS using anti-hu-IgG-PE, and binding of IgM in 10% NHS using and anti-hu-IgM-FITC by flow cytometry. Flow cytometry data are represented by geometric mean fluorescent intensity (gMFI) values of bacterial populations. (C) Inner membrane permeabilization of Kp209_CSTR in the presence of 1% NHS or NHS depleted using *E. coli* MG1655, Kp209 or Kp209_CSTR (Δ*E. coli* NHS, ΔKp209 NHS, and ΔKp209_CSTR NHS, respectively). ΔKp209_CSTR NHS was supplemented with physiological concentrations of IgG (+IgG; 1.25 mg/ml in 1% NHS) or IgM (+IgM; 15 μg/ml in 1% NHS) isolated from NHS. Bacteria were incubated at 37°C in the presence of 1 μM SYTOX green nucleic acid stain, and inner membrane permeabilization (SYTOX fluorescence intensity) was detected after 60 minutes. (D) IgM binding to Kp209 and Kp209_CSTR in IgM isolated from NHS, depleted using *E. coli* MG1655, Kp209 or Kp209_CSTR (Δ*E. coli* IgM, ΔKp209 IgM, and ΔKp209_CSTR IgM, respectively). Bacteria were incubated with depleted or non-depleted IgM (45 μg/ml) for 30 minutes at 4°C. Binding was detected by flow cytometry using anti-hu-IgM-FITC. Fluorescence was measured using flow cytometry. Flow cytometry data are represented by geometric mean fluorescent intensity (gMFI) values of bacterial populations. (E) Inner membrane permeabilization of Kp209_CSTR in the presence of 1% NHS depleted using Kp209_CSTR (ΔKp209_CSTR NHS), supplemented with physiological concentrations of IgM isolated from NHS (15 μg/ml in 1% NHS), depleted using *E. coli* MG1655, Kp209 or Kp209_CSTR (+Δ*E. coli* IgM, +ΔKp209 IgM, and +ΔKp209_CSTR IgM, respectively). Bacteria were incubated at 37°C in the presence of 1 μM SYTOX green nucleic acid stain, and inner membrane permeabilization (SYTOX fluorescence intensity) was detected after 60 minutes. (B-E) Data represent mean ± standard deviation of three independent experiments. Statistical analysis was performed using a paired one-way ANOVA with a Tukey’s multiple comparisons test on SYTOX fluorescence intensity (C&E) or Log10-transformed gMFI data (B&D). Significance is shown as *p≤0.05; ** p≤0.005; ***p≤0.005, ****p≤0.0005.

The observation that NHS contained Kp209_CSTR-specific IgM raised the question whether this IgM played a role in classical pathway activation. Therefore, we incubated Kp209_CSTR in Kp209_CSTR depleted NHS and observed that membrane permeabilization was reduced (fig. 4c). Membrane permeabilization of *E. coli* was not altered in Kp209_CSTR depleted NHS, indicating that the membrane permeabilizing potential was not affected by the depletion (fig. S2f). NHS depletion using Kp209 or *E. coli* did not affect complement-mediated membrane damage on Kp209_CSTR, indicating that components required for complement activation on Kp209_CSTR are still present (fig. 4c). This suggested that KP209_CSTR-specific IgM was responsible for complement-mediated membrane damage. To confirm the role of IgM we supplemented Kp209_CSTR depleted NHS with polyclonal IgG or IgM purified from NHS and monitored bacterial membrane permeabilization. Addition of polyclonal IgM fully restored membrane permeabilization on Kp209_CSTR, whereas polyclonal IgG did not (fig. 4c).

To verify that Kp209_CSTR-specific IgM was responsible for classical pathway activation, we used the antibody depletion technique to deplete polyclonal IgM isolated form NHS. Similar to NHS, depletion of polyclonal IgM with Kp209 reduced the IgM binding to Kp209_CSTR but again residual binding was observed, which confirmed the presence of Kp209_CSTR-specific IgM (fig. 4d). Depletion of polyclonal IgM using Kp209 or Kp209_CSTR did not affect binding to *E. coli* (fig. S2g). Next, we supplemented NHS depleted with Kp209_CSTR with the different preparations of IgM to study the effect on membrane permeabilization of Kp209_CSTR. Both IgM depleted with Kp209 or *E. coli* was able to restore membrane permeabilization on Kp209_CSTR, in contrast to IgM depleted with Kp209_CSTR (fig. 4e). This confirmed that Kp209_CSTR-specific IgM is crucial for antibody driven complement activation on Kp209_CSTR. In summary, we found that human serum contains IgM specific for Kp209_CSTR that induces MAC-mediated inner membrane damage.

### PhoQ of Kp209_CSTR is locked in an active state

Genetic comparisons between Kp209 and Kp209_CSTR has previously revealed that only the gene *phoQ* was altered in Kp209_CSTR (Janssen et al., 2021). This gene codes for PhoQ, the sensor histidine kinase of the PhoPQ two-component regulatory system, which is involved in gene regulation (Groisman, 2001). The finding that specific IgM molecules bound to Kp209_CSTR but not Kp209 suggested that mutations in *phoQ* led to the display of a novel epitope targeted by IgM. This might be explained by a difference in expression between Kp209 and Kp209_CSTR, or that the epitope could be shielded on Kp209 and therefore is not available. To aid the identification of the epitope recognized by the Kp209_CSTR-specific IgM, we studied the changes in gene expression in Kp209_CSTR compared to Kp209. To this end, the transcriptomes of both Kp209 and Kp209_CSTR were compared (fig. 5). We observed that 124 genes were significantly upregulated in Kp209_CSTR, whereas 60 genes were downregulated (Log_2_(FC)>1, P_adj_<0.05) (Table S1). As the complement activating IgM should binds to an epitope on the surface of Kp209_CSTR, we searched for upregulated genes involved in the production or modification of bacterial surface structures. Multiple genes encoding for fimbriae, outer membrane proteins, porins and efflux pumps were upregulated. Furthermore, expression of numerous genes involved in cell envelope modification were increased as well. However, because of the wide range of upregulated genes, it was not possible to pinpoint a particular surface structure as target of Kp209_CSTR-specific IgM. Next, we investigated the possibility of IgM epitope shielding on Kp209. To this end, we focused on the expression of genes involved in capsule and LPS O-antigen production, which are both known to be able to shield antibody targets on Gram-negative bacteria (Domínguez-Medina et al., 2020; Held et al., 2000). However, expression of neither capsule nor O-antigen genes was significantly altered in Kp209_CSTR compared to Kp209.

**Figure 5.**
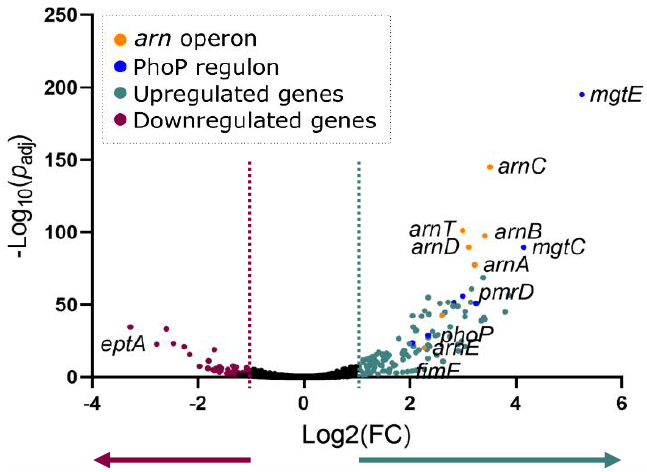
PhoQ of Kp209_CSTR is locked in a constitutively active state. RNA was isolated from log phase Kp209 and Kp209_CSTR, and their transcriptomes were analysed. The volcano plot depicts differential expressed genes in Kp209_CSTR compared to Kp209. In Kp209_CSTR 124 genes were upregulated and 60 downregulated.

Although we were not able to identify target of the Kp209_CSTR-specific IgM, we did observe that many of the upregulated genes in Kp209_CSTR were known to be upregulated after PhoQ activation. These include genes coding for proteins involved in magnesium transport, which is in line with the known magnesium sensory function of PhoQ, and *pmrD* which product activates PmrAB, another two-component regulatory system leading to LPS modifications (Groisman, 2001; Kato et al., 1999). The deletion found in Kp209_CSTR is within the sensory domain of PhoQ. The transcriptomic data indicates that this mutation leads to transcriptional activation of the PhoPQ regulon and suggests that PhoQ is locked in a constitutively active state in Kp209_CSTR.

### Inactivation of *phoQ* reverts colistin resistance and increases MAC-mediated membrane permeabilization of Kp209_CSTR

The constitutively active PhoQ influenced many genes that can alter the cell envelope composition and thereby MAC sensitivity. To verify whether PhoQ activity was required for MAC sensitivity, we screened a transposon (Tn) library of Kp209_CSTR for serum resistance. The library was first challenged with 16% NHS, followed by re-exposure of surviving bacteria to 32% NHS. Eight bacterial colonies that survived the challenges with 32% NHS were picked, and their *phoQ* genes were analysed. This revealed that three out of the eight Tn mutants had a transposon insertion in *phoQ*. Two of the three Tn∷*phoQ* mutants were clonally related as they had the same transposon barcode and identical insertion site. All Tn mutants showed a reduction in membrane permeabilization compared to Kp209_CSTR (fig. 6a), including the Tn∷*phoQ* mutants, confirming the involvement of PhoQ in MAC sensitivity. Since we observed that Kp209_CSTR-specific IgM was required for MAC induced membrane permeabilization, we analysed IgM binding to the Tn mutants. This revealed that Kp209_CSTR-specific IgM binding to the two unique Tn∷*phoQ* mutants was lost (fig. 6b). Interestingly, Kp209_CSTR-specific IgM binding was lost in most of the other Tn mutants as well, which emphasized the importance of IgM binding in MAC-driven membrane permeabilization. Kp209_CSTR-specific IgM binding was unaffected in one Tn mutant, indicating that loss of MAC-mediated permeabilization depended on an IgM-independent mechanism. Lastly, we investigated the colistin resistance of the Tn mutants. As colistin disrupts the bacterial cell envelop to kill bacteria, the membrane permeabilization assay was used to determine colistin sensitivity. Both Kp209_CSTR Tn∷*phoQ* mutants lost their colistin resistance due to the transposon insertion in *phoQ* (fig. 6c). Three MAC resistant Tn mutants remained colistin resistant, demonstrating that MAC resistance could occur in colistin resistant strains as well. Together these data further confirmed that colistin resistance and MAC sensitivity depended on PhoQ activity in Kp209_CSTR, since inactivation of PhoQ reverted colistin resistance and MAC-sensitivity.

**Figure 6.**
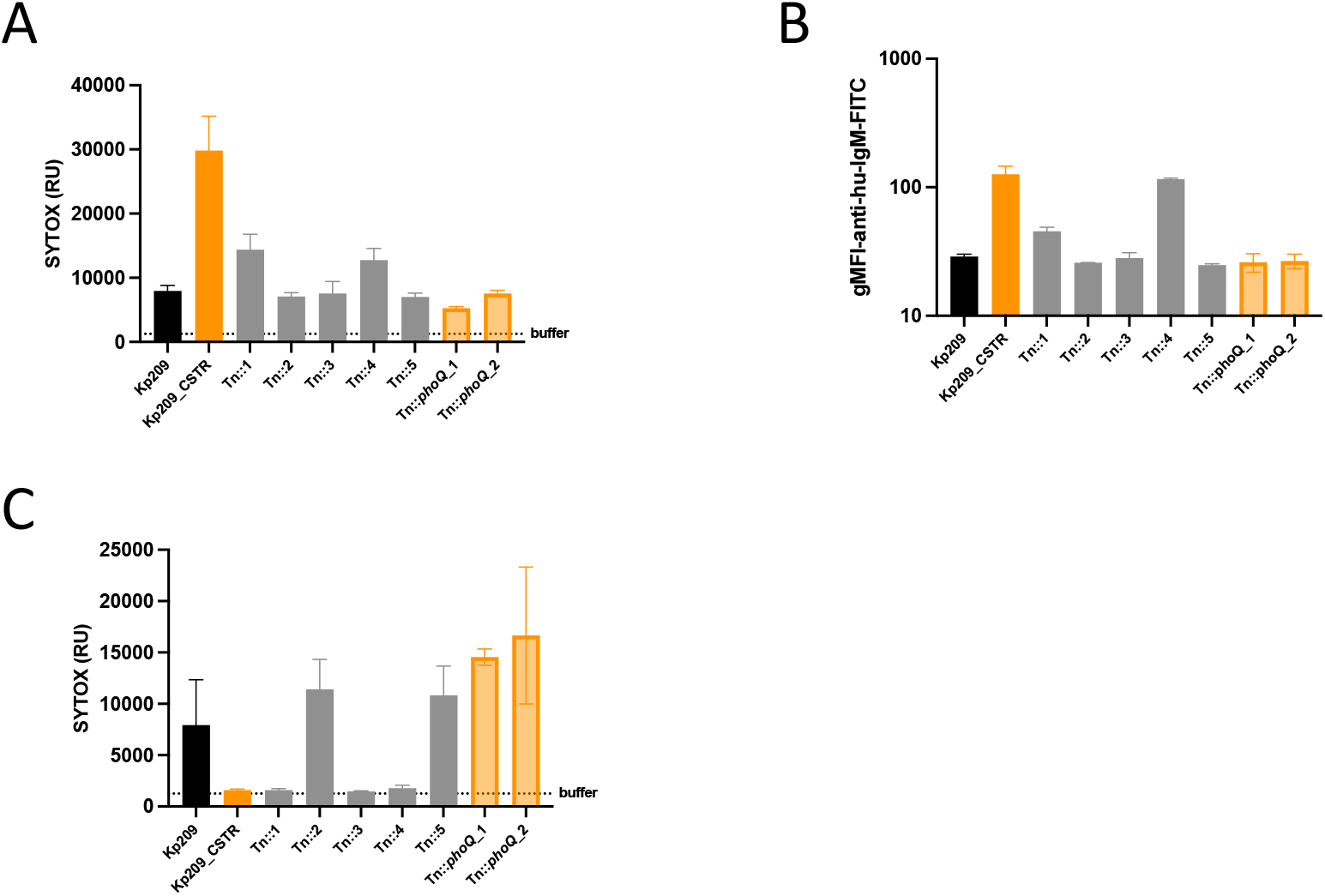
Transposon insertion in *phoQ* reverses the Kp209_CSTR phenotype back to Kp209. Kp209_CSTR transposon (Tn) library was exposed to 16% normal human serum (NHS), followed by re-exposed to 32% NHS, and surviving colonies were selected. Two unique Tn mutants had a transposon insertion in *phoQ* (Tn∷*phoQ* in translucent orange; other Tn mutants in grey). Kp209 and Kp209_CSTR are depicted in black and solid orange, respectively. (A) Inner membrane permeabilization of Kp209, Kp209_CSTR, and Kp209_CSTR Tn mutants treated with 10% NHS. Bacteria were incubated at 37°C in the presence of 1 μM SYTOX green nucleic acid stain, and inner membrane permeabilization (SYTOX fluorescence intensity) was detected after 60 minutes in a microplate reader. (B) IgM binding to Kp209, Kp209_CSTR, and Kp209_CSTR Tn mutants in 10% NHS depleted using, Kp209. Bacteria were incubated with Kp209 depleted or non-depleted NHS for 30 minutes at 4°C. IgM binding was determined using anti-hu-IgM-FITC by flow cytometry. Flow cytometry data are represented by geometric mean fluorescent intensity (gMFI) values of bacterial populations. (C) Inner membrane permeabilization of Kp209, Kp209_CSTR, and Kp209_CSTR Tn mutants in the presence of 1 μg/ml colistin. Bacteria were incubated at 37°C in the presence of 1 μM SYTOX green nucleic acid stain, and inner membrane permeabilization (SYTOX fluorescence intensity) was detected after 60 minutes in a microplate reader. (A-C) Data represent mean ± standard deviation of three independent experiments.

### Capsule production was reduced in Kp209_CSTR

Colonies of Kp209 appeared slightly more opaque than Kp209_CSTR, similar to what was observed between wild-type and capsule mutants in other *K. pneumoniae* strains (fig. S3a). Mutations in *phoQ* have previously been linked to altered capsule production in Gram-negative bacteria (Choi & Ko, 2015; Choi et al., 2020). Therefore, we decided to determine the production of capsule by quantifying the uronic acid content, an important component of the capsule (Clements et al., 2008). Although both Kp209 and Kp209_CSTR produced capsule, the production by Kp209_CSTR was reduced compared to Kp209 (fig. 7). Furthermore, we could show that this reduction was the result of the constitutively active PhoQ, as disrupting this gene by transposon insertion restored capsule production to level of Kp209 (fig. 7). The LPS O-antigen, the other major extracellular polysaccharide of *K. pneumoniae* did not differ in length and abundance between Kp209 and Kp209_CSTR when analysed by SDS-PAGE (fig. S3b). In summary, a slight change in capsule content was observed in Kp209_CSTR.

**Figure 7.**
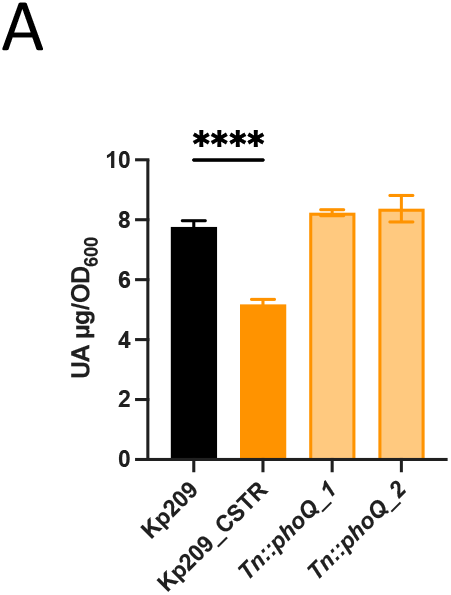
Kp209_CSTR has a reduced capsule production. Capsule production of Kp209 and Kp209_CSTR, and the Kp209_CSTR Tn∷*phoQ* mutants was determined by measuring uranic acid content. Absorbance was measured at 520nm, the uronic acid content calculated using a glucuronolactone standard curve and normalized to the culture density at 600nm. Data represent mean ± standard deviation of three independent experiments. Statistical analysis was performed using a paired one-way ANOVA with a Tukey’s multiple comparisons test. Significance is shown as ****p≤0.0005.

## Discussion

Membrane attack complex (MAC) insertion into the outer membrane is important for the direct killing of Gram-negative bacteria by the complement system. In the past, resistance against the membrane-targeting antibiotic colistin has been shown to influence sensitivity to other membrane-acting compounds in Gram-negative bacteria (Janssen et al., 2021; Napier et al., 2013). Here, we aimed to study the effect of developing colistin resistance on MAC-mediated killing of *K. pneumoniae*. We showed that colistin resistance mutations in *phoQ* led to increased MAC-mediated killing of the colistin resistant strain Kp209_CSTR. MAC-mediated membrane permeabilization in NHS was driven by IgM that specifically recognized Kp209_CSTR, indicating that the mutation in *phoQ* results in the exposure of a novel epitope on Kp209_CSTR. We observed that the mutation in *phoQ* locked PhoQ in an active state, which led to both increased colistin resistance, as well as increasing to MAC-mediated membrane permeabilization of Kp209_CSTR in NHS. *Vice versa*, inactivation of *phoQ* decreased MAC-dependent membrane permeabilization and colistin resistance.

Our mechanistic data demonstrates that MAC-mediated membrane permeabilization of Kp209_CSTR is driven by IgM specific for Kp209_CSTR in NHS. This suggests that an epitope for IgM becomes exposed on Kp209_CSTR due to the mutation in *phoQ*. The epitope might be shielded by capsular polysaccharides on the wild-type Kp209, and becomes available on Kp209_CSTR due to the reduction in capsule production. A role for a thicker capsule in shielding bacterial outer membrane and associated antigens from recognition by antibodies was demonstrated in previous studies. For example, chemical repression of capsule production by *K.* pneumoniae was demonstrated to lead to enhanced monoclonal antibody binding to the LPS (Domenico et al., 1999; Salo et al., 1995). Similarly, reduced capsule production due to genetic modifications changed the ability of the capsule to shield outer membrane proteins of *K. pneumoniae* (Held et al., 2000; Merino et al., 1992).

NHS and IgM was isolated from healthy donors with no described prior exposure to *K. pneumoniae*. The question remains why NHS contained IgM that specifically recognized Kp209_CSTR. One possibility is that the that IgM targeted Kp209_CSTR was a natural antibody, which are defined as germline-encoded antibodies expressed without any known direct antigenic stimulus (Holodick et al., 2017). Natural antibodies targeting bacterial antigens have been found in mice, and are assumed to exist in humans as well (Zhou et al., 2007). However, *K. pneumoniae* is a human commensal residing in the gut microbiota and nasopharynx, and estimations of gastrointestinal carriage in the western world range from 5 to 45% (Lepuschitz et al., 2020; Shimasaki et al., 2019). It is therefore probable that the serum IgM binding to Kp209_CSTR is part of a specific antibody response against antigens originating from commensal *K. pneumoniae.* This is supported by reports that healthy individuals produce both IgM and IgG antibodies recognizing *K. pneumoniae* (DeLeo et al., 2017; Garbett et al., 1989; Kobayashi et al., 2018; Lepper et al., 2003; Rossmann et al., 2015). Furthermore, it has been described that healthy donors possessed highly affinity-maturated antibodies belonging to the IgG, IgA and IgM isotypes that recognizing *K. pneumoniae* antigens (Pennini et al., 2017; Rollenske et al., 2018).

Our data indicates that an important determinant whether an IgM induces MAC-mediated killing is dependent on the recognized epitope. Only a minority of IgMs that bound to the surface of *K. pneumoniae* induced MAC-mediated membrane permeabilization in Kp209_CSTR. A large part of the IgM that bound Kp209_CSTR also bound to Kp209, but these IgM were not involved in inducing MAC-mediated inner membrane damage. However, we showed that only Kp209_CSTR-specific IgM was crucial for inner membrane permeabilization. The difference between these IgM indicate that the IgM target is an important determinant for MAC-dependent membrane damaging effects. This difference might relate to the location of the target in relation to the outer membrane. For *K. pneumoniae* it has been reported that MAC deposition needs to occur close to the outer to be bactericidal, and that localization of MAC far from the outer membrane prevents bacterial lysis (Jensen et al., 2020). This would suggest that the target of the bactericidal IgM would be located in close proximity to the outer membrane. Further studies might elucidate the what surface structure is targeted by the IgM but both LPS and outer membrane proteins remain potential targets (Domenico et al., 1999; Serushago et al., 1989; Xin Wang-Lin et al., 2017).

We found that colistin resistance mutations in *phoQ* had an opposite effect on the MAC sensitivity of Kp209_CSTR. This might be explained by the different modes of action between colistin and MAC. Although colistin and MAC both interact with and destabilize bacterial membranes, their mechanisms are critically different. Colistin is a small amphipathic molecule attracted electrostatically to the negatively charged cell envelope, where it inserts into and destabilizes the bacterial membranes (Andrade et al., 2020). Antibody-induced MAC deposition is initiated after specific recognition of bacterial surface structures by antibodies. This activates of the classical complement cascade, resulting in sequential deposition of complement components on the bacterial surface. Only in the final stages of the complement cascade, when C5 convertases are being deposited, can membrane piercing MAC pores can be formed (Heesterbeek et al., 2019). Localization of C5 convertases on the bacterial surface is crucial for MAC-dependent membrane permeabilization, as well as proper C9 polymerization (Doorduijn et al., 2021; Heesterbeek et al., 2019). As colistin and MAC target bacteria by different mechanisms, becoming resistant to colistin could have a different effect on MAC sensitivity. In contrast to what we observe for Kp209_CSTR, colistin resistance did not enhance MAC sensitivity of Kp257_CSTR and Kp040_CSTR under the tested conditions. This indicates that colistin resistance can occur without influencing MAC sensitivity, which is supported by the observation that several Kp209_CSTR Tn mutants were resistant to both MAC and colistin. Colistin resistance due to mutations in *phoQ* has been linked serum sensitivity in both *K. pneumoniae* and *E. coli* (Choi & Ko, 2015; Choi et al., 2020). It was shown that colistin resistant mutants were more serum resistant compared to wild-type strains. These colistin resistant mutants had mutations in *phoQ,* modifications in lipid A, as well as a reduced capsule production (Choi & Ko, 2015; Choi et al., 2020). Our work adds to the growing understanding how the PhoPQ two-component regulation system influences both colistin and serum sensitivity.

Transcriptome analysis indicated that PhoQ was more active in Kp209_CSTR compared to Kp209. The *phoQ* gene of Kp209_CSTR contains a WAQRN-97-C deletion that is located in the sensory domain of PhoQ (Janssen et al., 2021; Lemmin et al., 2013). In *Salmonella enterica* the tryptophan residue at position 97 (W-97) in PhoQ has been reported to be crucial for magnesium sensing, and loss of PhoQ’s sensory function leads to a constitutively active form of PhoQ (Chamnongpol et al., 2003; García Véscovi & Soncini, 1996). In concordance with our own results, mutations in the PhoQ sensory domain in *K. pneumoniae* ST23 have been reported to lead to upregulation of PhoPQ regulated genes, indicating that PhoQ was more active (Choi & Ko, 2015). We found that PhoQ activity influences capsule production, which might play a role in binding of Kp209_CSTR-specific IgM. Transcriptomic analysis revealed that expression of capsule genes was not altered, suggesting that all the capsule producing machinery should be present in Kp209_CSTR. This suggests that capsule production was not affected on a transcriptional level, but at a later stage of capsule synthesis. It has been postulated that addition of 4-amino-4-deoxy-L-arabinose (L-Ara4N) to lipid A, one of the modifications observed for Kp209_CSTR, could influence capsule production (Grangeasse et al., 2003; Janssen et al., 2021). The precursor of L-Ara4N, UDP-glucuronic acid, is also required for capsule production (Grangeasse et al., 2003; Rodrigues et al., 2020; Wang et al., 2020). Increased L-Ara4N synthesis would limit the availability of UDP-glucuronic acid, thereby reducing capsule production. This is supported by reports that capsule production can be enhanced by mutating LPS syntheses genes in *E. coli* (Wang et al., 2020). UDP-glucuronic acid is converted to L-Ara4N by the Arn pathway. In line with the hypothesis, the *arnABCDET* operon genes are among the strongest upregulated genes in Kp209_CSTR. The *arn* operon is under control the PmrAB two-component system, which is activated by PmrD (Dalebroux & Miller, 2014). The expression of *pmrD* is regulated by PhoPQ and was strongly upregulated in Kp209_CSTR as a result of the mutations in *phoQ* (Groisman, 2001).

As the number of antibiotic resistant *K. pneumoniae* strains rises, infections with these bacteria becomes an increasing risk to human health. Studying how *K. pneumoniae* is killed by, and evades, the complement system will reveal new insights in the infection biology of *K. pneumoniae*. Understanding the interaction between killing of *K. pneumoniae* by the immune system and antibiotics will help to improve treatment of *K. pneumoniae* infections

## Materials & methods

### Bacterial strains

*Klebsiella pneumoniae* clinical isolates were collected during routine diagnostics in the medical microbiology department in the University Medical Centre Utrecht, The Netherlands. *Klebsiella pneumoniae* Kp209, Kp257 and Kp040, and their colistin resistant daughter strains were kindly provided by Axel Janssen (University Medical Centre Utrecht, The Netherlands; University of Lausanne, Switzerland;(Janssen et al., 2021)). KPPR1S and KPPR1S*ΔwcaJ* were kindly provided by Kimberly Walker (University of North Carolina, USA;(Walker et al., 2020)). For the experiments with *E. coli* the laboratory strain MG1655 was used.

### Serum preparation and reagents

Normal human serum (NHS) was prepared as described before (Heesterbeek et al., 2019). In short, blood was drawn from healthy volunteers, allowed to clot, and centrifuged to separate serum from the cellular fraction. Serum of 15-20 donors was pooled and stored at −80°C. Heat inactivation (Hi) of NHS was achieved by incubating NHS at 56°C for 30 minutes. OMCI was produced and purified as previously described (Nunn et al., 2005). RPMI (ThermoFisher) supplemented with 0.05% human serum albumin (HSA, Sanquin), further referred to as RPMI buffer, was used in all experiments, unless otherwise stated. Eculizumab was kindly provided by Frank Beurskens, Genmab, Utrecht, The Netherlands. Sera deficient from factor B (fB) and complement component C2 were obtained from Complement Technology. Purified factor B and C2 were produced by U-protein express (Utrecht, The Netherlands). C1r protease inhibitor BBK32 was kindly provided by Brandon Garcia, Greenville, NC, USA. Monoclonal mouse anti-hu-C1q IgG1 4A4B11 was produced in house (ATCC HB-8327) (Reckel et al., 1986; Zwarthoff et al., 2021).

### Bacterial growth

For all experiments, bacteria were cultured on Lysogeny broth (LB) 1.5% agar plates at 37 °C, unless stated otherwise. Single colonies were picked and cultured overnight in liquid LB medium at 37 °C while shaking. The following day the bacteria were subcultured by diluting the overnight culture 1/100 in fresh medium and grown to OD_600_= 0.4-0.5 at 37 °C while shaking. Bacteria were washed twice with RPMI buffer by centrifugation at 10,000 g for two minutes and resuspended to OD_600_= 0.5 in RPMI buffer. For the Kp209_CSTR Tn mutants, both solid and liquid medium were supplemented with 30 μg/ml kanamycin.

### Bacterial viability assay

Bacteria (OD_600_= 0.05) were incubated in serum diluted in RPMI buffer. In the conditions where C5 conversion was blocked to prevent the initiation to the terminal complement pathway, 20 μg/ml OMCI + 20 μg/ml Eculizumab was added. After incubation at 37 °C, 10-fold serial dilutions in RPMI buffer were prepared and plated. Colony forming units were determined on LB agar plates after culturing the bacteria on plate over night at 37 °C. Survival in NHS was normalized to NHS in which C5 conversion was inhibited ((#NHS/#C5 inhibition) *100%).

### Membrane permeabilization

Bacteria (OD_600_= 0.05) were incubated in serum or colistin diluted in RPMI buffer in the presence of 1 μM SYTOX Green Nucleic Acid stain (ThermoFisher). Bacteria were incubated at 37 °C under shaking conditions. Fluorescence was determined in a microplate reader (CLARIOstar, Labtech) using an excitation wavelength of 490-14 nm and a emission wavelength of 537-30 nm.

### Antibody deposition

Bacteria (OD_600_= 0.05) were incubated in HiNHS diluted in RPMI buffer for 30 minutes at 4 °C under shaking conditions. Bacteria were washed twice by centrifugation with RPMI buffer, resuspended 1 μg/ml goat anti-hu-IgG-AF647 (2040-31, SouthernBiotech) or 2 μg/ml goat anti-hu-IgM-FITC (2020-02, SouthernBiotech) for 30 minutes at 4 °C while shaking. Bacteria were washed twice with RPMI buffer and fixated in 1.5% paraformaldehyde in PBS for five minutes. Fluorescence was determined via flow cytometry (MACSQuant, Milteny Biotech), acquiring 10,000 events per condition. Flow cytometry data was analysed in FlowJo V.10. Bacteria were gated based on the forward and side scatter, and AF647 and FITC geometric mean fluorescence intensity was determined for the bacterial population.

### Serum depletion with bacterium

Bacteria (OD_600_= 1.0) were incubated in ice cold NHS (20% in RPMI buffer). After a 10-minute incubation on ice while shaking, bacteria were pelleted in a cooled centrifuge (two minutes at 10.000 g) and the bacterium depleted NHS was collected. The depletion steps were preformed trice in total. Antibody depletion was verified via flow cytometry.

### Polyclonal IgG and polyclonal IgM isolation from NHS

IgG and IgM were isolated from NHS as previously described (Zwarthoff et al., 2021). In short, IgG was isolated using 5 ml HiTrap Protein G High Performance column (GE Healthcare), whereas IgM was isolated using POROS™ CaptureSelect™ IgM Affinity Matrix (ThermoScientific) in a XK column (GE Healtcare) using an ÄKTA FPLC system. After capture antibodies were eluted according to the manufacturer’s instructions. Collected antibodies were dialyzed overnight against PBS at 4 °C, and stored at −80 °C.

### LPS silver stain

LPS silver stain was performed as previously described (Doorduijn et al., 2021). Briefly, single bacterial colonies were scraped of agar plates, and heat-inactivated at 56 °C for one hour, followed by protein digestion using proteinase K (400 μg/ml) for 90 minutes at 60 °C. Samples were diluted in 2x Laemmli buffer with 0.7 M β-mercaptoethanol, ran over a 4-12% BisTris gel and fixed overnight in ethanol (40% v/v) + glacial acetic acid (4% v/v). The gel was oxidised for five minutes in ethanol (40% v/v) + glacial acetic acid (4% v/v) + periodic acid (0.6% m/v) and stained for fifteen minutes in silver nitrate (0.6% m/v in 0.125 M sodium hydroxide + 0.3% ammonium hydroxide v/v). The gel was developed for seven minutes in citric acid (0.25% m/v) + formaldehyde (0.2% v/v).

### Capsule production analysis

Capsule production was assessed by measuring the uronic acid content as previously described (Walker et al., 2020). Briefly, 500 μl stationary phase bacteria grown in LB medium added to 100 μl 1% Zwittergent 3-14 detergent in citric acid (100 mM pH 2.0) and incubated at 50 °C for 20 minutes. 300 μl supernatant was collected after centrifugation (five minutes 16,000 g), mixed with 1200 μl 100% ethanol, and incubated at 4 °C overnight. After centrifugation (five minutes 16,000 g), the pellet was dissolved in 200 μl distilled water and 1200 μl tetraborate (12.5 mM in sulfuric acid) was mixed in by vortexing. After a five-minute incubation at 100 °C, the samples were cooled and 20 μl 3-hydroxydiphenol (0.15% in 0.5% sodium hydroxide) was added. Absorbance was measured at 520 nm. Uronic acid amounts were calculated from a standard curve prepared with glucuronolactone and normalized to the culture density (OD_600_).

### Transposon library and serum exposure

Kp209_CSTR was mutagenized via conjugation using strain WM3064 carrying the pKMW7 vector with a barcoded Tn5 transposon library as described previously (Wetmore et al., 2015). Bacteria with a transposon insertion were selected using LB supplemented with kanamycin and the library was stored (OD_600_= 1) at −80 °C. For serum exposure experiments, transposon library was 10-fold diluted in LB medium and cultured to mid-log (OD_600_= 0.6). Bacteria were pelleted and resuspended in RPMI buffer. Washed bacteria (OD_600_= 0.05) were incubated with RPMI buffer or 16% NHS for 2 hours at 37 °C. Samples were diluted 10-fold in LB medium and incubated overnight at 37 °C. The overnight culture was diluted 100-fold in fresh LB medium and challenged with 32% NHS for 2 hours at 37 °C followed by serial dilutions in PBS and plating on LB agar. Surviving colonies were picked that survived the 32% NHS challenge for further analysis.

### Transcriptome analysis

#### RNA extraction

Subcultures of Kp209 and Kp209_CSTR were grown to OD_600_=0.5-0.6 at 37 °C while shaking. RNA was isolated using the hot phenol-chloroform method. Briefly, bacterial were incubated in hot phenol lysis solution (1% SDS, 2 mM EDTA, 40 mM sodium acetate in acid phenol (Invitrogen #15594-047)) for 45 minutes at 65 °C. Total RNA was purified by successive steps of phenol-chloroform extraction, followed by an extraction with cold chloroform. After ethanol precipitation, genomic DNA was digested by Turbo DNAse (Invitrogen #AM1907) treatment. Total RNA quantity and quality were assessed using an Agilent Bioanalyzer.

#### RNA-seq libraries construction

Ribosomal RNAs (rRNAs) depletion was performed using Ribominus bacteria 2.0 transcriptome isolation kit (Invitrogen) according to manufacturer’s instructions. Depletion efficacy was verified on an Agilent Bioanalyzer using RNA pico chips and remaining RNAs were concentrated using ethanol precipitation. The Collibri stranded RNA library prep kit for Illumina (Invitrogen) was used to build cDNA libraries for sequencing starting from 15 ng of RNA according to manufacturer’s instructions. cDNA libraries quality and concentration were assessed using an Agilent Bioanalyzer High sensitivity DNA chip. RNA-seq was performed in biological triplicates.

#### Sequencing and data analysis

Sequencing was performed by the Montpellier GenomiX platform (MGX, https://www.mgx.cnrs.fr/) using an Illumina Novaseq instrument. Following quality control, Bowtie2 was used for sequencing reads mapping (Langmead & Salzberg, 2012; Seemann, 2014) onto the genome of Kp209 available on the European Nucleotide Archive under accession number PRJEB29521 (Janssen et al., 2021), previously annotated with Prokka. The read count per feature was calculated using Ht-seq count (Anders et al., 2015). Finally, differential gene expression between the Kp209 and Kp209_CSTR strains was performed using DESeq2 (Love et al., 2014).

## Data analysis and statistical testing

Unless stated otherwise data collected as three biological replicates and analysed using GraphPad Prism version 9.4.1 (458). Statistical analyses are further specified in the figure legends.

## Data availability

The RNA-seq data generated in this study were deposited on the NCBI Gene Expression Omnibus (GEO) under the GSE212413 accession number

## Ethics statement

Human blood was isolated after informed consent was obtained from all subjects in accordance with the Declaration of Helsinki. Approval was obtained from the medical ethics committee of the UMC Utrecht, The Netherlands.

## Acknowledgments

This work was supported by the Netherlands Organization for Scientific Research (NWO) through a TTW-NACTAR Grant #16442 (to SvdL and SHMR) and the European Union’s Horizon 2020 research programs 787 H2020-EU-ITN-EJD (CORVOS #860044 to F.M. and SHMR). The authors would like to thank Frank Beurskens (Genmab) for providing Eculizumab, and Brandon Garcia for providing BBK32.

---

**Figure S1.**
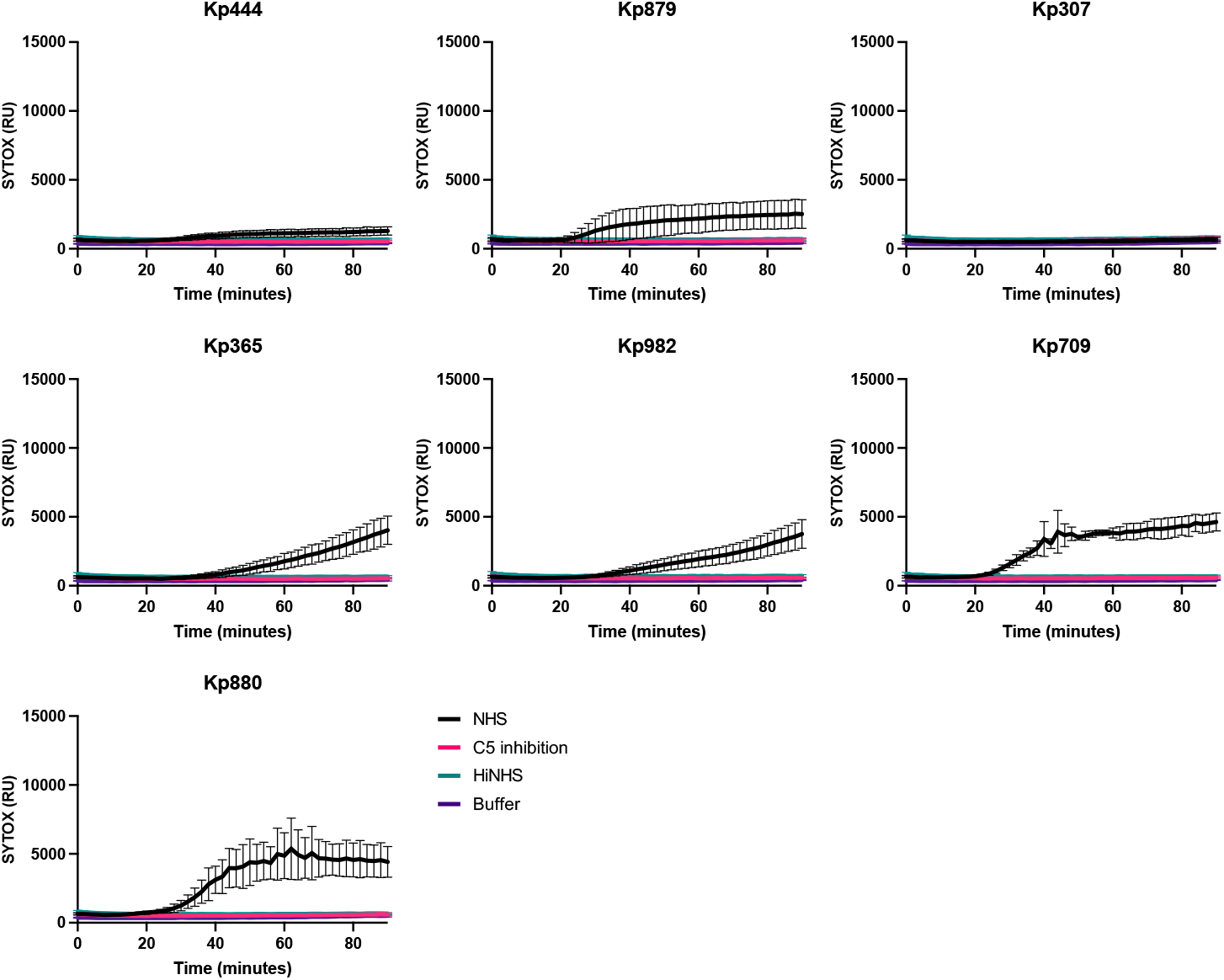
Inner membrane permeabilization of Kp444, Kp879, Kp307, Kp365, Kp982, Kp709 and Kp880 in time. Bacteria were incubated in the presence of 10% NHS, 10% NHS in which C5 conversion was inhibited by addition of 20 μg/ml OMCI and 20 μg/ml Eculizumab (C5 inhibition), or 10% heat inactivated NHS (HiNHS), at 37°C in the presence of 1 μM SYTOX green nucleic acid stain, and inner membrane permeabilization (SYTOX fluorescence intensity) was detected every 2 minutes for 90 minutes in a microplate reader. Data represent mean ± standard deviation of three independent experiments.

**Supplemental figure 2.**
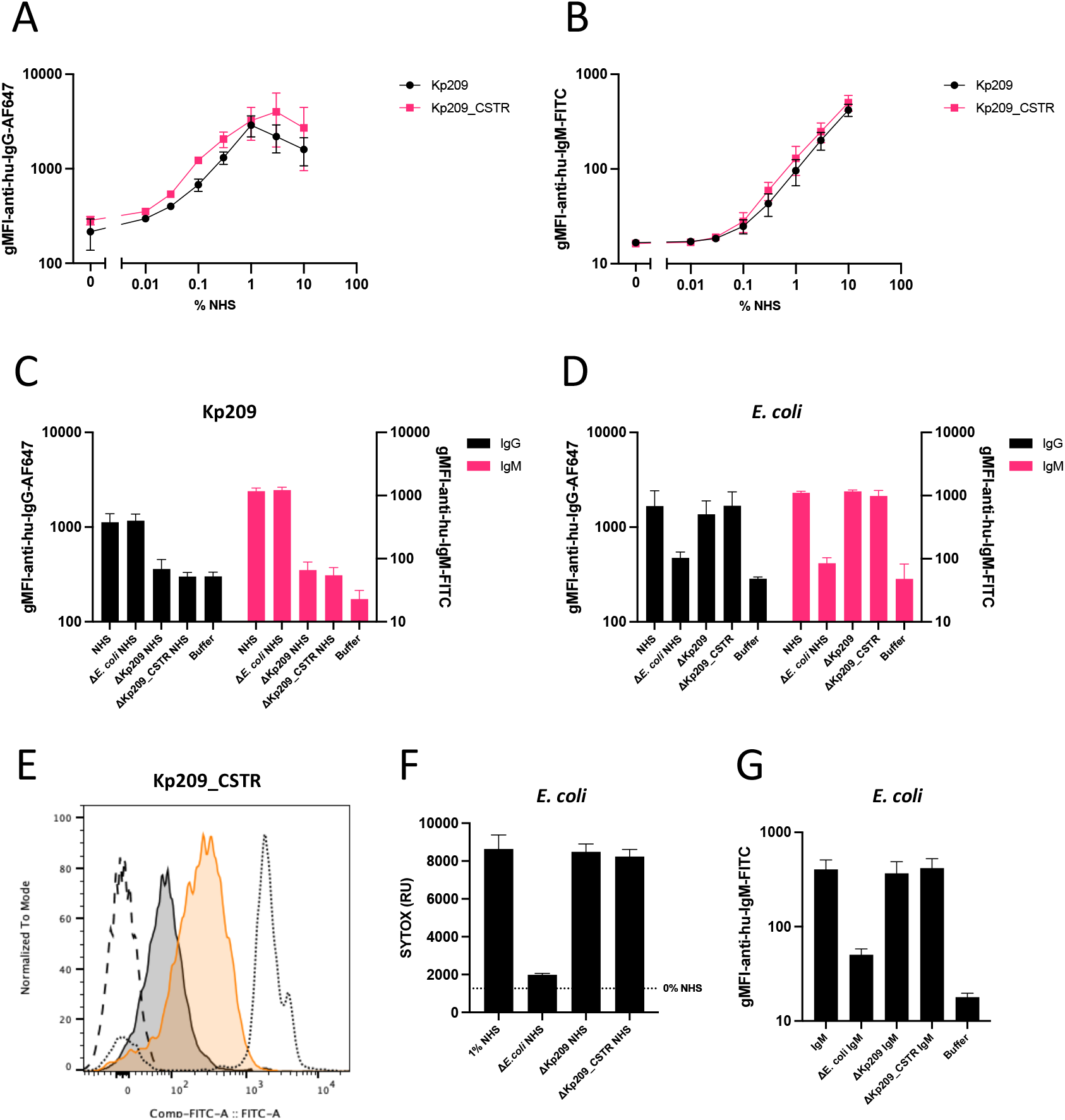
(A&B) Antibody binding to Kp209 and Kp209_CSTR in NHS. Bacteria were incubated with NHS for 30 minutes at 4°C. Total IgG (A) and total IgM (B) binding was determined using anti-hu-IgG-PE and anti-hu-IgM-FITC, respectively, by flow cytometry. Flow cytometry data are represented by geometric mean fluorescent intensity (gMFI) values of bacterial populations. (C&D) Antibody binding to Kp209 (C) and *E. coli* MG1655 (D) in in NHS depleted using *E. coli* MG1655, Kp209 or Kp209_CSTR (Δ*E. coli* NHS, ΔKp209 NHS, and ΔKp209_CSTR NHS, respectively). Bacteria were incubated with depleted or non-depleted NHS for 30 minutes at 4°C. Binding of IgG was measured in 0.3% NHS using anti-hu-IgG-PE, and binding of IgM in 10% NHS using anti-hu-IgM-FITC by flow cytometry. Flow cytometry data are represented by geometric mean fluorescent intensity (gMFI) values of bacterial populations. (E) IgM binding to Kp209 and Kp209_CSTR with IgM (45 μg/ml) isolated from NHS (dotted), IgM depleted with Kp209 (grey) or Kp209_CSTR (orange). Kp209_CSTR was incubated with depleted or non-depleted IgM (45 μg/ml) for 30 minutes at 4°C, and binding was detected via flow cytometry using and anti-hu-IgM-FITC. Slitted line corresponds to the buffer control. Geometric mean fluorescent intensity (gMFI) and number of events normalized to mode are depicted on the X- and Y-axis, respectively. The graph is representative for three independent repeats. (F) Inner membrane permeabilization of *E. coli* MG1655 in the presence of 1% NHS, or NHS depleted using *E. coli* MG1655, Kp209 or Kp209_CSTR (Δ*E. coli* NHS, ΔKp209 NHS, and ΔKp209_CSTR NHS, respectively). Bacteria were incubated at 37°C in the presence of 1 μM SYTOX green nucleic acid stain, and inner membrane permeabilization (SYTOX fluorescence intensity) was detected after 60 minutes. (G) IgM binding to *E. coli* MG1655 using IgM isolated from NHS, depleted using *E. coli* MG1655, Kp209 or Kp209_CSTR (Δ*E. coli* IgM, ΔKp209 IgM, and ΔKp209_CSTR IgM, respectively). Bacteria were incubated with depleted or non-depleted IgM (45 μg/ml) for 30 minutes at 4°C. Binding of IgM was detected using anti-hu-IgM-FITC by flow cytometry. Flow cytometry data are represented by geometric mean fluorescent intensity (gMFI) values of bacterial populations. (A-D&F-G) Data represent mean ± standard deviation of three independent experiments. Statistical analysis was performed using a paired two-way ANOVA with a Šidák multiple comparisons test on Log10-transformed gMFI data (A&B), or a paired one-way ANOVA with a Tukey’s multiple comparisons test on SYTOX fluorescence intensity (F) or Log10-transformed gMFI data (C, D&G).

**Figure S3.**
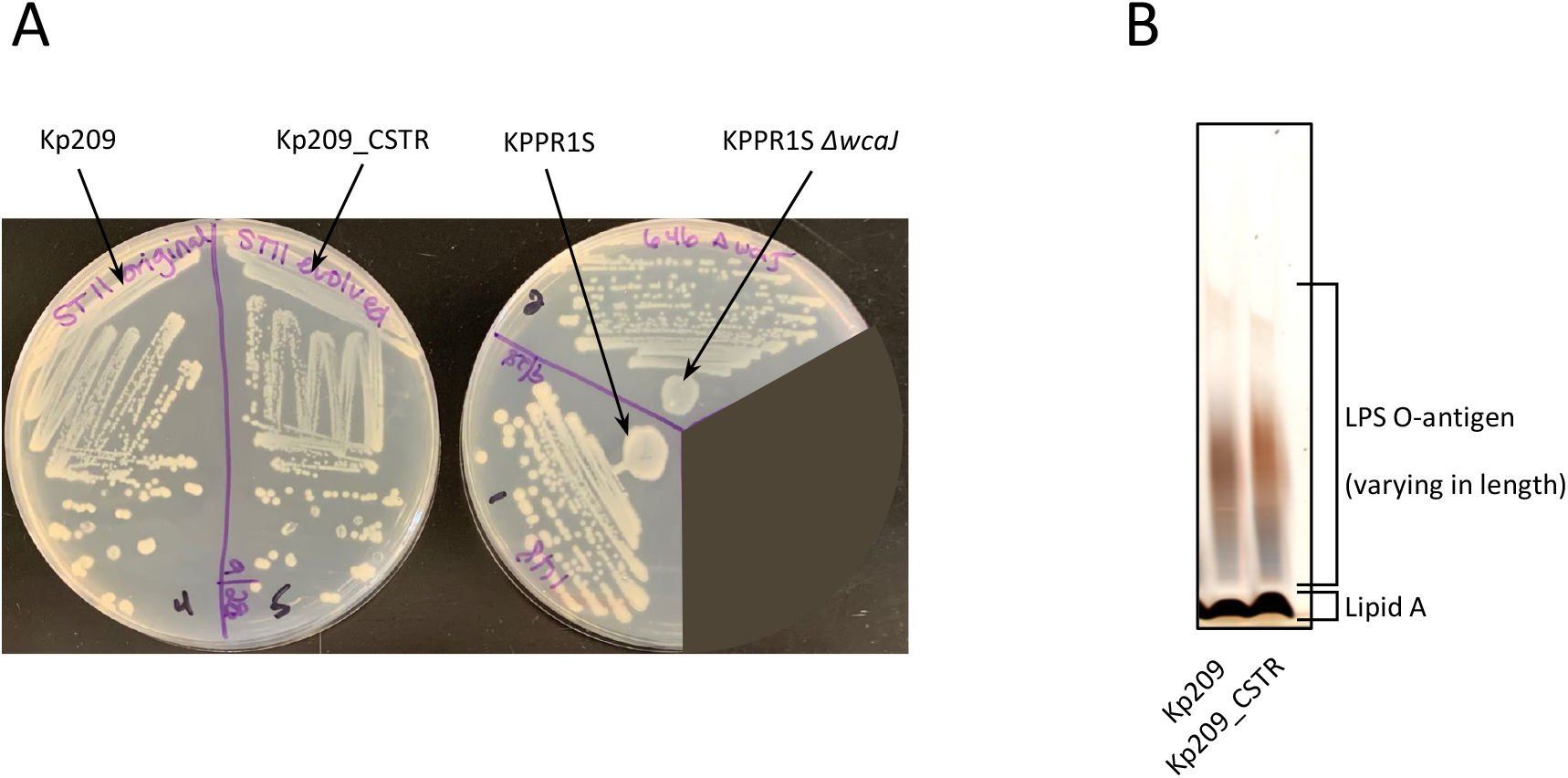
(A) Image of Kp209, Kp209_CSTR, KPPR1S and capsule mutant KPPR1S *ΔwcaJ* cultured overnight on LB agar plates. Both Kp209_CSTR and KPPR1S *ΔwcaJ* appear more opaque on plate compared to their parental strains Kp209 and KPPR1S, respectively. (B) SDS-PAGE silver stain of Kp209 and Kp209_CSTR LPS. Bacterial lysates were digested with protein K to remove all proteins, and loaded on a SDS-PAGE gel. The LPS lipid A could, based on size, be distinguished from the O-antigen, which can have various lengths. Image is a representative of two independent experiments.

**Table S1.**
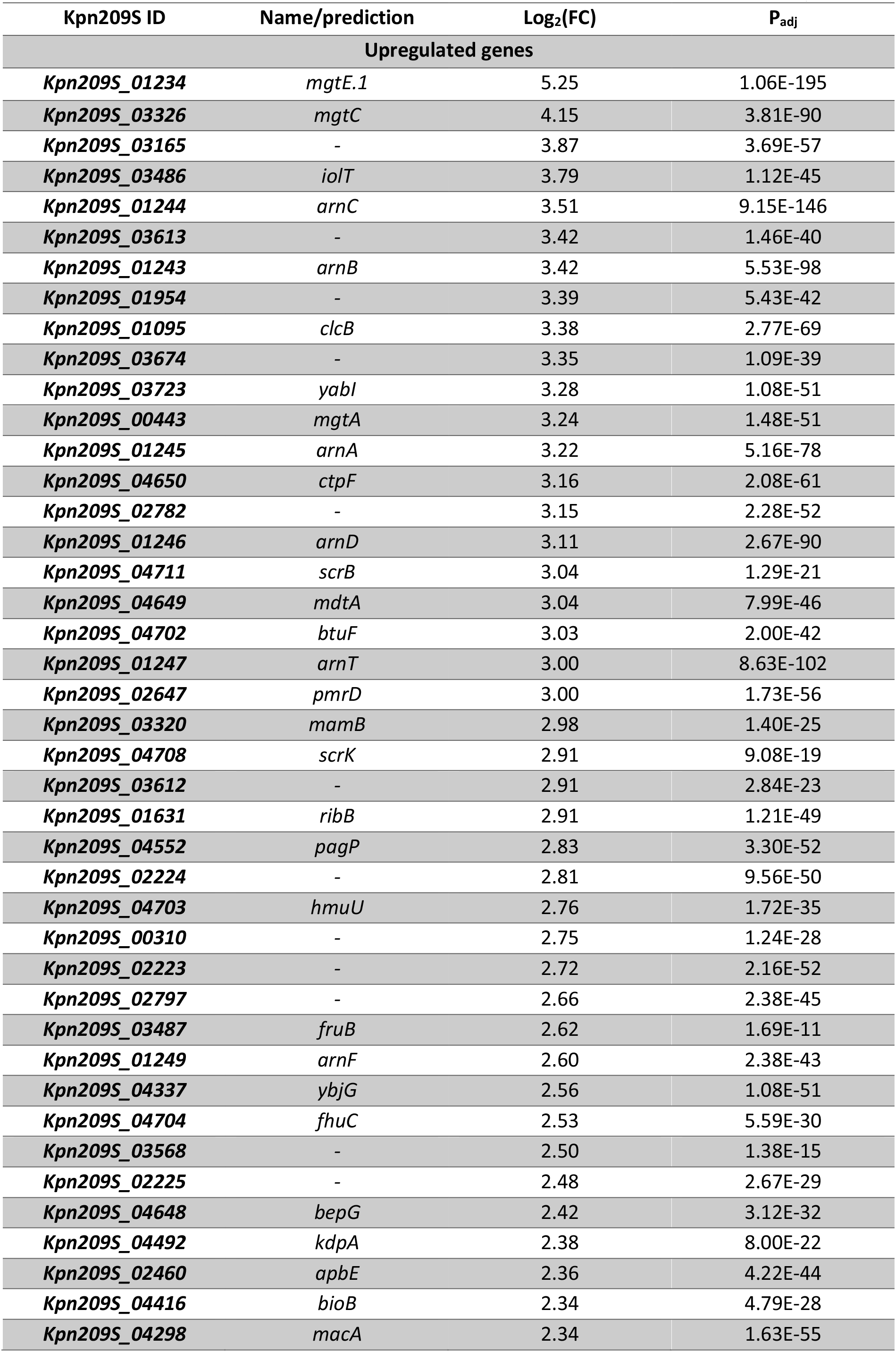
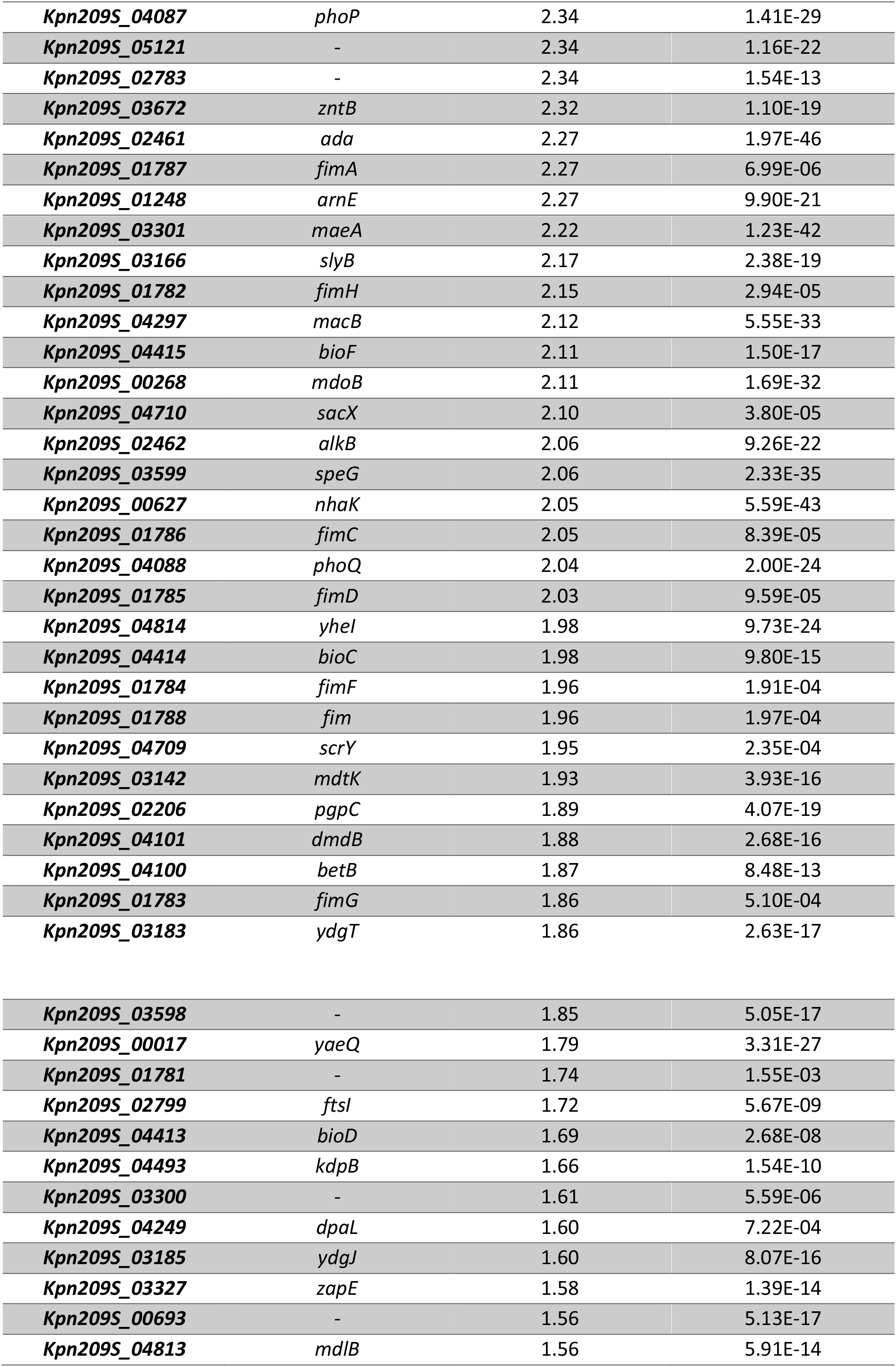
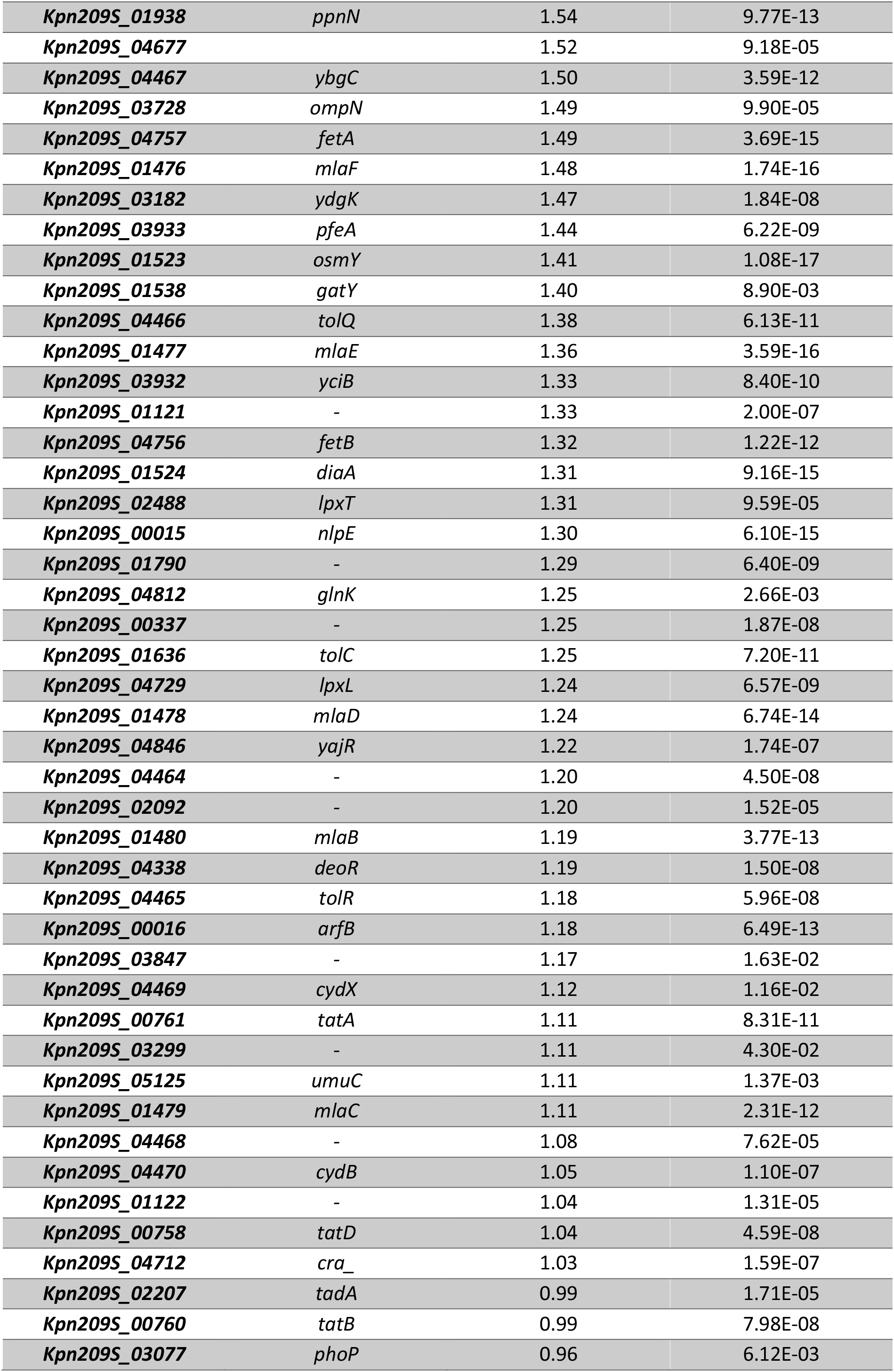
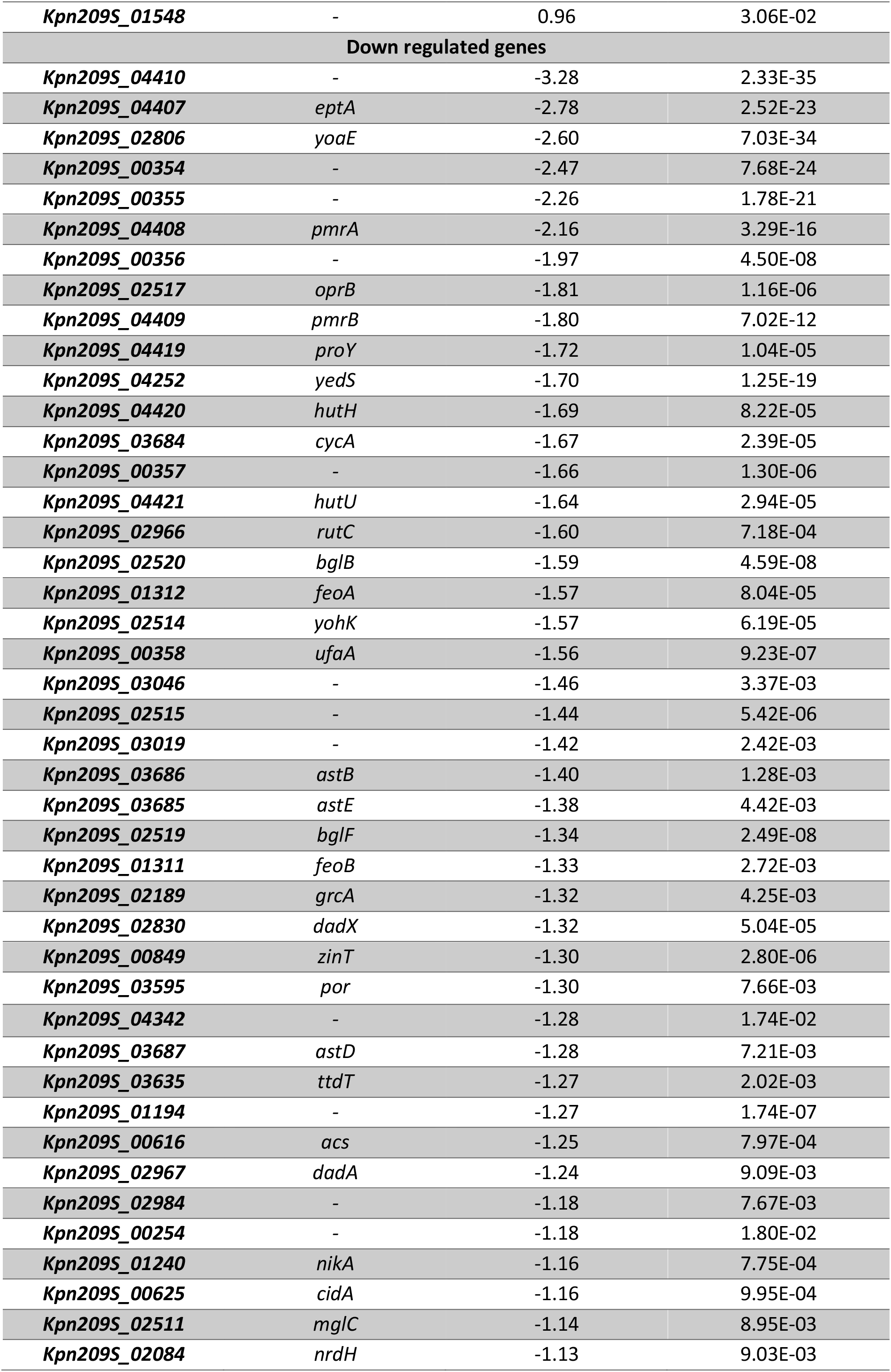
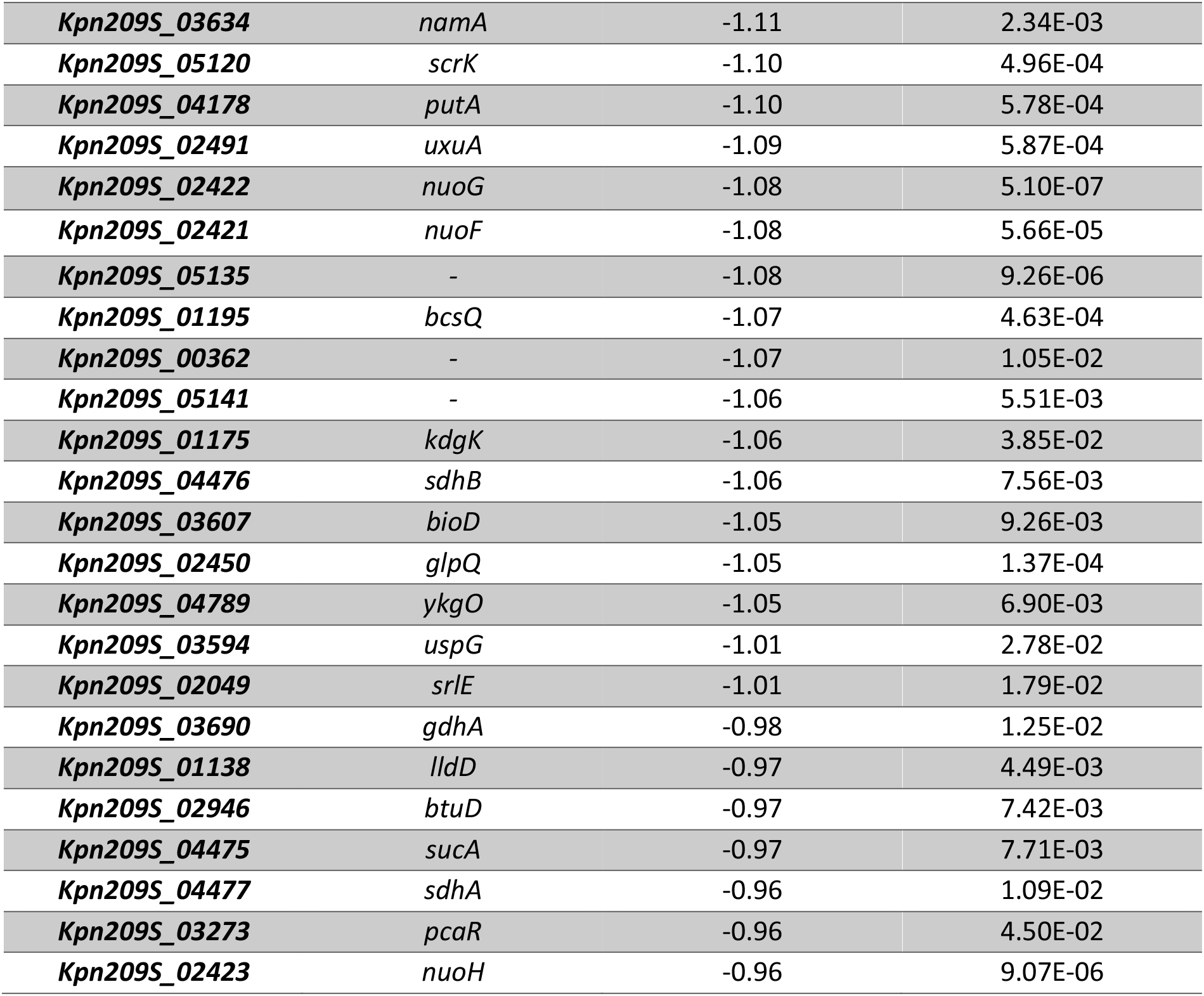
Differentially expressed genes in Kp209_CSTR compared to Kp209 (Log_2_(FC)>1, P_adj_<0.05)

## Notes

### Competing Interest Statement

The authors have declared no competing interest.

